# High-Throughput Antibody Neutralization Screening in Massively Parallel Droplet Arrays

**DOI:** 10.1101/2025.01.31.635965

**Authors:** Matias Gutierrez-Gonzalez, Ahmed S. Fahad, Cyrille L. Delley, Cheng-Yu Chung, Shuyan Jin, Nicoleen Boyle, Matheus Oliveira de Souza, Azady Pirhanov, Nicolas M.S. Galvez, Camila T. França, Samantha Marglous, Evan Bhagat, Donovan Vincent, Daniel Neumeier, Yi Cao, Nicole Doria-Rose, Sai T. Reddy, Aaron G. Schmidt, Alejandro B. Balazs, Adam R. Abate, Brandon J. DeKosky

**Affiliations:** The Ragon Institute of Mass General, MIT, and Harvard, Cambridge, MA, USA; Department of Chemical Engineering, Massachusetts Institute of Technology, Cambridge, MA, USA; Department of Bioengineering and Therapeutic Sciences, University of California San Francisco, San Francisco, CA, USA; Department of Pharmaceutical Chemistry, The University of Kansas, Lawrence, KS, USA; Department of Biosystems Science and Engineering, ETH Zurich, Basel, Switzerland; Vaccine Research Center, National Institute of Allergy and Infectious Diseases (NIAID), National Institutes of Health (NIH), Bethesda, MD, USA; Department of Microbiology, Harvard Medical School, Boston, MA, United States; Koch Institute for Cancer Research, Cambridge, MA, USA; Department of Chemical Engineering, The University of Kansas, Lawrence, KS, USA

## Abstract

Neutralizing antibodies provide rapid immune defense against infectious diseases, but are difficult to discover at scale because neutralization assays require live reporter cells and soluble monoclonal antibodies. Here we report Droplet Reporter Cell Testing for Neutralization (DrReCT-Neutralization) to screen antibody gene libraries for their ability to neutralize viral infections. We established the necessary engineered cell lines and validated the DrReCT screening platform using synthetic oligoclonal libraries, followed by an example discovery campaign that demonstrated scalable functional antibody data collection against viral diseases.

## Main Text

Neutralizing antibodies are defined by their ability to suppress viral infection *in vitro*, and neutralizing antibody titers are closely correlated with vaccine protection for several pathogens^1^. The study of neutralizing monoclonal antibodies (mAb) has also become a major focus for vaccine design^2^. Despite the increasing importance of neutralizing mAb discovery, the current antibody discovery technologies cannot efficiently screen for neutralization directly. Most large-scale (*i.e.*, >1,000 variant) discovery platforms screen cellular or protein display libraries based on binding affinity to soluble protein antigens; example technologies include phage^3^ or yeast^4–7^, and the B cell receptor (BCR) displayed on the surface of memory B cells. Affinity-based discovery relies on the availability of engineered soluble extracellular domains that can be difficult (or impossible) to produce, as many viral receptors are incompletely characterized or cannot be purified in a natural, membrane-bound form. Affinity-based screening also cannot account for alternate receptors and co-receptors^10^. Some receptor blocking assays can pre-enrich for antibodies against known neutralization-vulnerable epitopes^8^, but viruses present multiple vulnerable epitopes and binding affinity to a non-native soluble viral protein does not directly correlate with neutralizing activity. Evaluating antibody neutralization therefore still requires follow-up live cell assays using soluble antibody protein in well plates, where capital- and reagent-intensive robotics are necessary to achieve more than 500 tests within a few days^9^.

In addition to antigen production and affinity screening difficulties, screening full-length antibodies in therapeutically relevant formats (e.g., IgG1) has proven challenging. Single-cell microfluidic assays using secreted animal-derived B cells showed the potential of droplet screening based on binding affinity, but these affinity-based workflows still rely on purified antigen proteins and cannot screen for critical functional properties like neutralization^11–13^. One group reported progress with multi-cell in-droplet, on-chip microfluidic neutralization sorts^14^. However, the technical requirements for multi-cell microfluidics in-line with on-chip fluorescence detection & droplet sorting are very high. Furthermore, platform validation was restricted to low-diversity clonal spike-ins without quantitative analyses of a cell library by next-generation sequencing (NGS) or the discovery of new clones. Thus there is currently no scalable platform to screen diverse, full-length soluble mAb libraries directly for neutralization function.

To overcome these limitations, we established a high-throughput approach to analyze mAb gene libraries via microfluidic encapsulation of antibody-secreting reporter cells, followed by cell recovery for efficient fluorescence-activated cell sorting (FACS) and sensitive NGS quantitation of the sorted libraries (**Fig. 1**). In the DrReCT-Neutralization assay, antibody gene libraries are engineered to secrete from reporter cell lines. Next, we use a microfluidic cell encapsulation system to isolate single antibody-secreting cells in stable emulsions at a rate of 10^6^ cells per hour. Secreted antibodies accumulate inside droplets, and subsequent microdroplet picoinjection introduces a pseudovirus challenge to the droplets (**Fig. 1A**). Infection inside droplets leads to reporter gene expression, and cell libraries are recovered for conventional FACS to separate infected and uninfected cells. Finally, NGS quantitatively identifies the mAbs that were enriched in uninfected cells and were depleted in infected cells – the neutralizing antibodies – which are prioritized for recombinant production and follow-up traditional assays. The DrReCT-Neutralization platform is also highly flexible in nature and is compatible with a variety of infectious disease models by simply changing the reporter cell lines and the virus/pseudovirus challenge.

**Figure 1.**
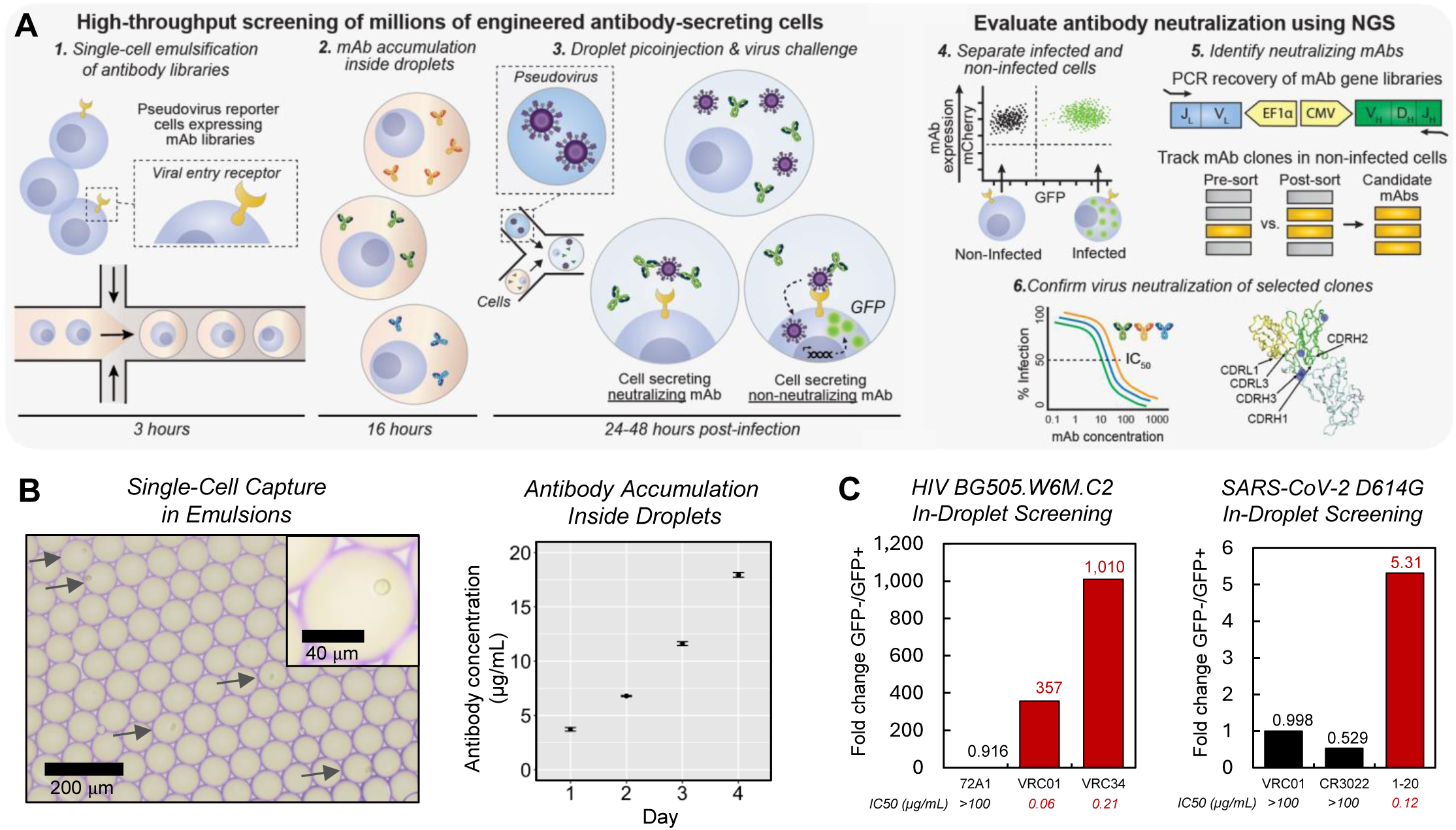
Establishment of DrReCT-Neutralization, a flexible platform for massively parallel antibody neutralization screening using droplet arrays. **A.** A library of antibody-secreting cells is isolated by single-cell emulsification. After sufficient antibody accumulation inside droplets, the cells in each droplet are challenged with a reporter virus or pseudovirus. Infected cells express the reporter gene (GFP shown here), and cells are recovered from droplets and fractionated by FACS into GFP+ (infected) and GFP− (uninfected) populations. Antibody gene libraries are amplified and analyzed using sensitive NGS. mAb clonal prevalence is compared between the infected and non-infected groups to reveal clones with improved neutralization function. **B.** Secretion of a control antibody 1-20 by HEK/ACE2 cells isolated inside droplets was quantified by ELISA over 4 days. Error bars indicate standard deviation of the mean for three technical replicates. **C.** Synthetic oligoclonal pools were screened as an example experiment using two different pseudovirus reporter challenge models (HIV-1 and SARS-CoV-2). Cells expressing neutralizing antibodies were spiked into a background of control cells expressing non-neutralizing antibodies. Oligoclonal pools were then analyzed with the DrReCT-Neutralization screening platform to test platform effectiveness for real-world applications. Antibody read frequencies between GFP+ and GFP− sorted cell populations were used to calculate antibody clonal fold-change values. Compiled data for Panel C is provided in **Table S1**.

To establish a proof-of-concept and establish initial parameters for the assay, we first tested performance in 96-well plates with antibody-secreting reporter cell lines challenged with a SARS-CoV-2 D614G pseudovirus encoding green fluorescent protein (GFP) (**Fig. S1**). HEK/ACE2 cells were engineered to secrete either a known neutralizing antibody (910-30 or 1-20, with IC50 values of 0.13 and 0.12 μg/mL, respectively), or a non-neutralizing antibody (VRC01 or CR3022). After transfection, single cells were seeded in 96-well plates, and the secreted antibody was allowed to accumulate for 12 days until cell density reached confluence prior to pseudovirus challenge. As expected, following pseudovirus challenge the cells secreting neutralizing antibodies had lower GFP reporter expression than non-neutralizing controls (**Fig. S1**). We also separately verified performance of the picoinjection system using a tracking dye, which enables precise temporal control of the in-droplet pseudovirus challenge (**Fig. S2**).

We next evaluated the capacity of engineered antibody-secreting reporter cells to survive and secrete antibody at sufficient concentrations inside droplets to exceed reasonable neutralizing IC50 values (**Fig. 1B**). HEK/ACE2 cells expressing the SARS-CoV-2 neutralizing antibody 1-20 (IC50 = 0.12 μg/mL) were encapsulated inside droplets, cultured over the course of several days, and recovered at various time points to quantify in-droplet antibody concentrations. We found that cells effectively secreted antibodies for at least four days, and concentration increased linearly from 4 μg/mL after one day to 16 μg/mL after four days. Thus by adjusting droplet incubation time prior to virus/pseudovirus challenge (via picoinjection), we reasoned that our assay can detect neutralizing antibodies with an IC50 up to approximately 20 μg/mL with the current engineered antibody secretion and incubation methods.

Next, we performed two demonstration campaigns to quantify assay performance with synthetically generated oligoclonal libraries. We spiked a small number of engineered cells secreting known neutralizing mAbs into background populations of cells secreting non-neutralizers. As positive controls, we used HIV-1 neutralizing antibodies (VRC01 and VRC34) and a SARS-CoV-2 neutralizer (1-20). We sought to detect quantitative differences in the prevalence of neutralizing antibodies in the infected (GFP+) versus non-infected (GFP−) populations after droplet-based challenge with either the HIV-1 (BG505.W6M.C2) or SARS-CoV-2 (D614G) pseudovirus strains (**Fig. 1C**, **Table S1**). After libraries were screened as shown in **Figure 1A**, we calculated the frequency of each mAb gene in infected versus non-infected populations. NGS analysis revealed that neutralizing mAbs were effectively depleted from infected (GFP+) populations, and were reliably detected based on a high GFP−/GFP+ prevalence ratio in sorted libraries. Using this metric, the quantitative signal for neutralizing clones exceeded 300-fold for the high-throughput HIV-1 neutralization assay, and 5-fold for the SARS-CoV-2 assay, after only a single round of screening (**Table S1**). In this experiment the HIV-1 pseudovirus infected ~30% of cells inside droplets, whereas SARS-CoV-2 pseudovirus infected ~1% of cells inside droplets; the stronger enrichment signals in HIV-1 assays was likely due to more efficient HIV-1 in-droplet infection. These data demonstrate that quantitative signals from neutralizing antibodies can still be detected with in-droplet infection efficiencies less than 5%, still, we increased the SARS-CoV-2 pseudovirus concentration in later experiments to enhance quantitative assay metrics and cell throughput.

After validating DrReCT-Neutralization with synthetic oligoclonal mixtures, we next conducted an antibody engineering campaign to identify mutations that improve neutralization against an evolved SARS-CoV-2 strain, XBB.1.16, that was first identified in January 2023. We started from the neutralizing antibody S2E12 (first reported in 2020)^15^ and performed site-saturation mutagenesis (SSM) to create a gene library with all possible single amino acid substitutions at each position of the heavy chain variable region (**Fig. 2A**). The SSM library was cloned into a lentiviral antibody expression plasmid containing an mCherry expression marker, packed into lentiviral particles, and transduced into HEK/ACE2 cells. Next, antibody-secreting cell libraries were co-captured in droplets along with XBB.1.16 GFP reporter pseudovirus, followed by cell recovery and sorting into GFP− and GFP+ populations for NGS analysis. After in-droplet pseudovirus challenge and cell recovery from emulsions, 23% of the recovered cells expressed both mCherry and the GFP infection reporter gene (**Fig. 2B**). NGS analysis revealed 1,083 single mutation S2E12 variants in screening data, and we selected 5 variants with high GFP−/GFP+ ratios and high prevalence in the pre-sort library (≥1 in 3,000 prevalence) for soluble IgG expression and traditional neutralization assays (**Fig. 2C**). We found that all 5 of the tested S2E12 variants had improved neutralization potency, ranging from 1.5-fold to 5.1-fold compared to the template antibody (**Fig. 2D**, **Fig. S3**, **Table S2**).

**Figure 2.**
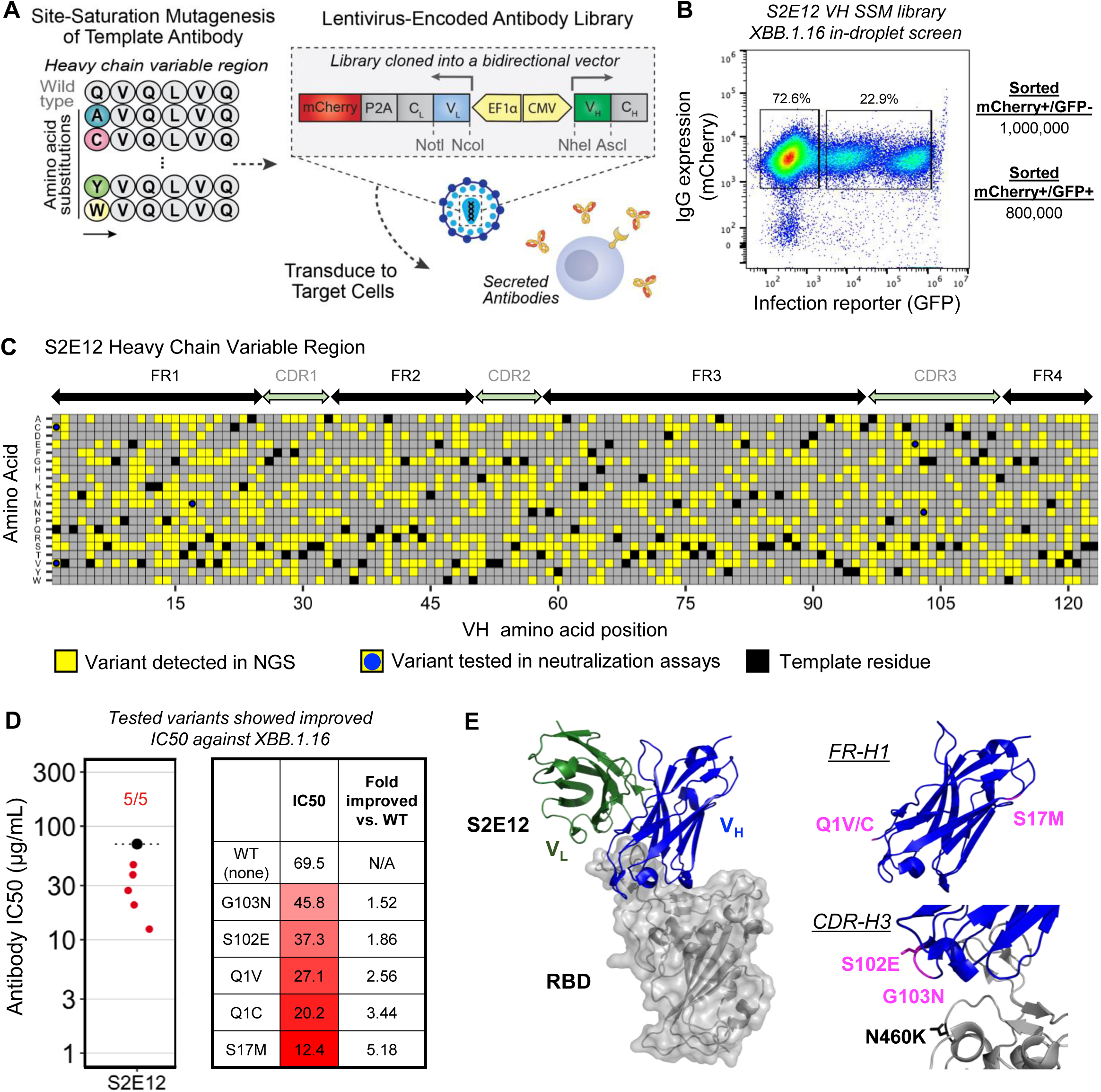
Discovery of improved neutralizing antibody variants from a site saturation mutagenesis (SSM) library. **A.** The heavy chain of S2E12 was diversified using SSM, packed into lentiviral particles, and the resulting library was transduced into HEK/ACE2 cells for high-throughput analysis by DrReCT-Neutralization. **B.** mCherry+/GFP− and mCherry+/GFP+ populations were sorted after recovery from in-droplet pseudovirus challenge; recovered cell counts are shown on the right. Over 1 million sorted cells were recovered from the high-throughput single-cell assay. **C.** Map of high-throughput screening results from DrReCT-Neutralization, where infected and non-infected populations were sequenced after droplet challenge. Variants with coverage >0.85 are shown here, and the top 5 variants with the highest GFP−/GFP+ ratio were selected for soluble IgG expression and functional evaluation in traditional well-plate assays. Coverage was estimated based on transduction efficiency at the lentiviral library generation step. **D.** Plate-based neutralization assay results for expressed S2E12 variants against SARS-CoV-2 XBB.1.16. **E.** Mutations were mapped onto an S2E12 co-crystal structure with SARS-CoV-2 RBD to reveal structural locations of enhanced neutralization.

We mapped the locations of functionally-improving mutations onto the known S2E12 co-crystal structure with the SARS-CoV-2 receptor binding domain (RBD) to assess potential mechanism(s) of neutralization improvement (PDBs 7K3Q, 7K45). Two heavy chain mutations, S102E and G103N, were located in the CDR-H3 and may promote binding interactions with K460 in XBB1.16 through additional salt bridge or hydrogen bonding, respectively. However, the CDR-H3 in the known co-crystal structure is ~10-15 Å from K460, and would require repositioning to support these interactions. A co-crystal structure of the XBB1.16 variant in complex with S2E12 may show additional global repositioning of the VH and VL that may help facilitate these interactions. The other three functionally-improving mutations (Q1C/V a S17M) are located in the FR-H1 and represent the most significant improvements (~2.6 – 5.2 fold more potent). However, it is not readily apparent from the SARS-CoV-2 RBD:S2E12 complex how these mutations influence interactions with XBB1.16. One possible explanation is that allosteric effects modulate long-range interactions by repositioning CDRs in the antigen combining site. We note that traditional methods for designed mutagenesis would not normally identify mutations far from the binding site, which were most effective here. In fact, the Q1 residue was not resolved in some cryo-EM structures (e.g., PDB 7K45, 7K4N), demonstrating the power of high-throughput functional screening to identify beneficial mutations that would be difficult to predict by modeling known structures.

Our study applied DrReCT-Neutralization, a single-cell antibody secretion platform integrated with droplet microfluidics, to interrogate large populations of antibody-secreting cells based on neutralization function. In contrast to traditional protein affinity-based discovery methods, DrReCT-Neutralization screens for function against the native form of membrane-expressed viral receptors and does not require soluble/purified viral antigens for antibody discovery, nor does it require two different cell lines (one secreter, one reporter) to be isolated in a droplet and detected & sorted on-chip. DrReCT uses engineered cells that can accommodate any synthetic or natural protein gene library, including natively paired heavy:light configurations, and it screens for antibody function in a soluble, druggable-ready IgG format. These important simplifications from prior techniques improve screening throughput and quality while reducing cost. We note that during the discovery phase of this work, we never used robotic platforms nor expressed & purified a soluble viral receptor antigen, which both represent major bottlenecks for studies of recently emerged pathogens, and for viruses and evolved strains that have no reliable stabilized receptor antigens available.

Our ability to repeatedly screen cellular secretion libraries also supports functional analysis against multiple and expanded viral strains, which is critical to study mAbs against evolving pathogens like HIV-1, influenza, and SARS-CoV-2. Here we used lentiviral transduction to generate libraries, which does limit library size and can result in multiple transgene insertions and/or variable expression across cells. Future work with DrReCT could use targeted genomic integration tools to generate screening libraries, such as CRISPR/Cas9 or recombinases, for efficient single-site gene integrations^16^. We note that highly expressed mAb variants are more efficiently surveyed by DrReCT; future studies could bin libraries by antibody reporter gene expression to improve screening of poorly expressed variants. DrReCT is easily scalable to >10 million cells per experiment and the technology supports a wide range of studies, including functional data collection for AI/ML model training. Overall, DrReCT-Neutralization offers a powerful approach that expands the scale of antibody neutralization data collection and enables protein engineering based directly on neutralization function.

## Acknowledgements

We gratefully acknowledge M. Waring and I. Midtbust-Heger for assistance with flow cytometry, A. Antonelli for assistance with experiments, and D. Gludish and D. Russell for providing TZM-gfp cells.

## Author contributions

M.G.G., A.S.F., C.C., and B.J.D. conceived the study. M.G.G., A.S.F., C.C., N.D.R., S.R.R., A.B.B., A.R.A., and B.J.D. designed the experiments. M.G.G., A.S.F., C.C., S.J., N.B., A.P., N.M.S.G., S.M., Y.C., and A.G.S. conducted experiments or analyzed the data with the assistance of C.L.D., M.O.S., C.T.F, E.B., D.V., and D.N., S.T.R., A.B.B., A.R.A., and B.J.D. supervised the experiments. M.G.G., A.S.F., and B.J.D. wrote the manuscript with input from all authors. All authors contributed to the article and approved the submitted version.

## Conflict of Interest

Massachusetts Institute of Technology, Massachusetts General Hospital, and The University of Kansas have filed for patent protection on the technology described herein, and B.J.D., M.G.G., C.C. A.S.F, N.B., and S.J. are named as co-inventors.

## Data Availability Statement

Raw MiSeq sequencing data is deposited in the NCBI Short Read Archive (SRA) under accession number PRJNA1184180.

## Funding

This work was supported by NIH grants DP5OD023118, R01AI141452, R21AI143407, R21AI166396, U01AI169587, and 1R01AI181684, by the MIT Research Support Committee, the MIT Department of Chemical Engineering, and by the Ragon Institute of MGH, MIT, and Harvard. A.B.B. was supported by NIAID R01s AI174875, AI174276, the NIDA Avenir New Innovator Award DP2DA040254, the NIDA Avant-Garde Award 1DP1DA060607, a Massachusetts Consortium on Pathogenesis Readiness (MassCPR) grant, CDC subcontract 200-2016-91773-T.O.2 and a grant from Coalition for Epidemic Preparedness Innovations (CEPI).

## Supplementary Figures

**Figure S1.**
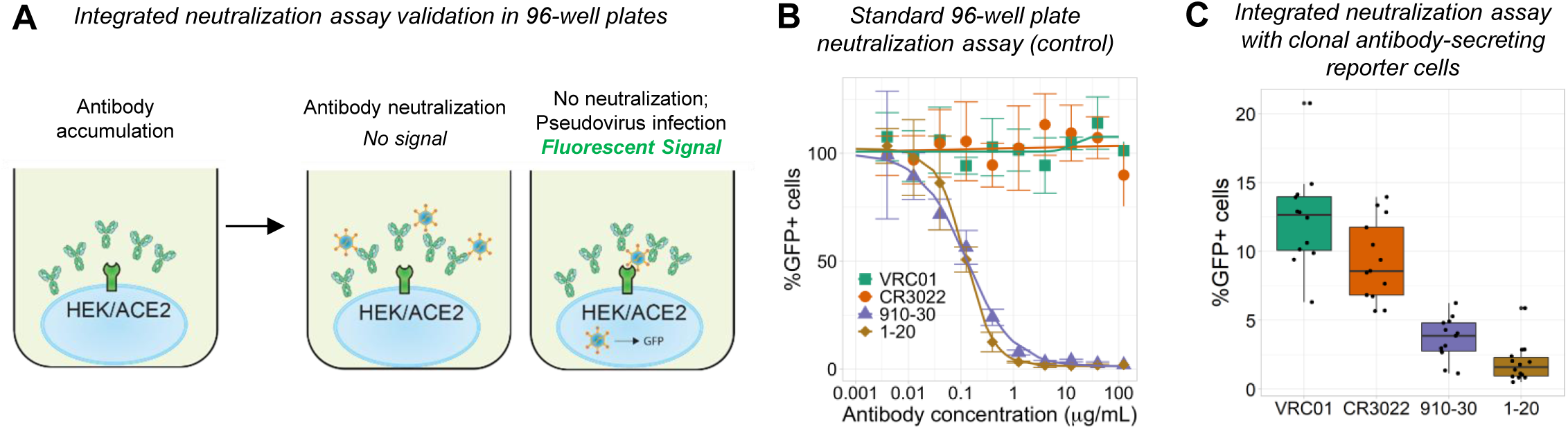
An integrated neutralization assay with both antibody secretion and viral challenge in 96-well plates. **A.** Overview of an integrated neutralization assay used to evaluate the feasibility of droplet neutralization tests. HEK/ACE2 cells expressing monoclonal antibodies are seeded in 96-well plates, and antibody accumulation occurs prior to the addition of SARS-CoV-2 reporter viral particles. **B.** A traditional 96-well plate neutralization assay using soluble IgG. Error bars indicate the standard deviation of the mean for n=3 replicates at each concentration. GFP expression was measured by flow cytometry. **C.** An integrated neutralization assay in 96-well plates. First HEK/ACE2 cells were transfected with plasmids encoding for different monoclonal antibodies and clonal populations were selected by limiting dilution and antibiotic selection. Next, single cells were seeded into the wells, and neutralization tests were carried out as indicated in Panel A.

**Figure S2.**
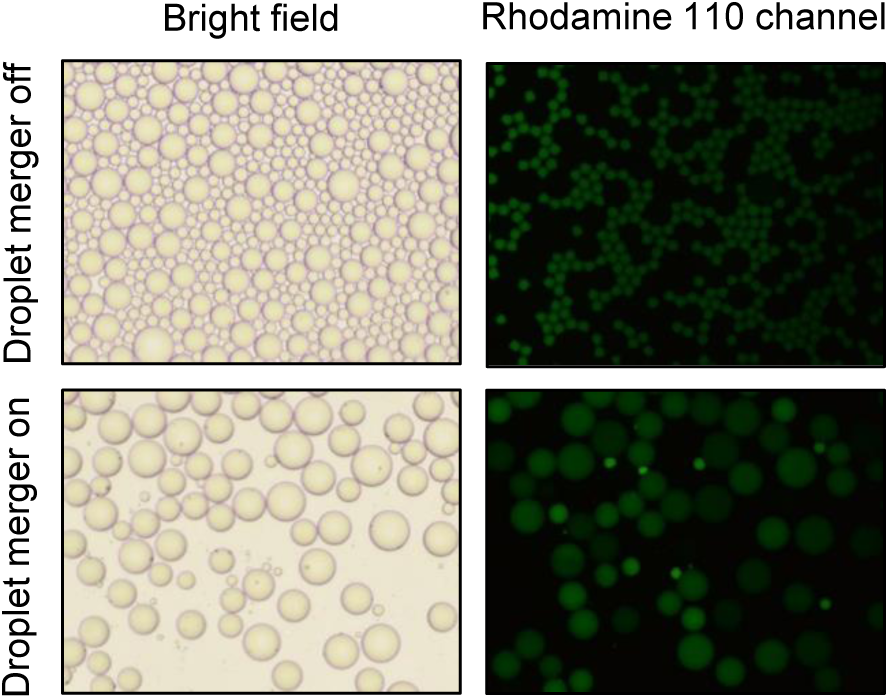
Pico-injection method validation. ~80 μm diameter emulsion droplets containing PBS were generated and injected into a custom microfluidic device designed for pico-injection^17^. Next, new droplets ~40 m in diameter containing rhodamine 110 were generated and pico-injected (merged) with the PBS droplets when passing through an electric field. The output of the device was collected with the electrical field off and on, and pictures were captured in both bright field and a rhodamine 110 channel to confirm appropriate droplet merging.

**Figure S3.**
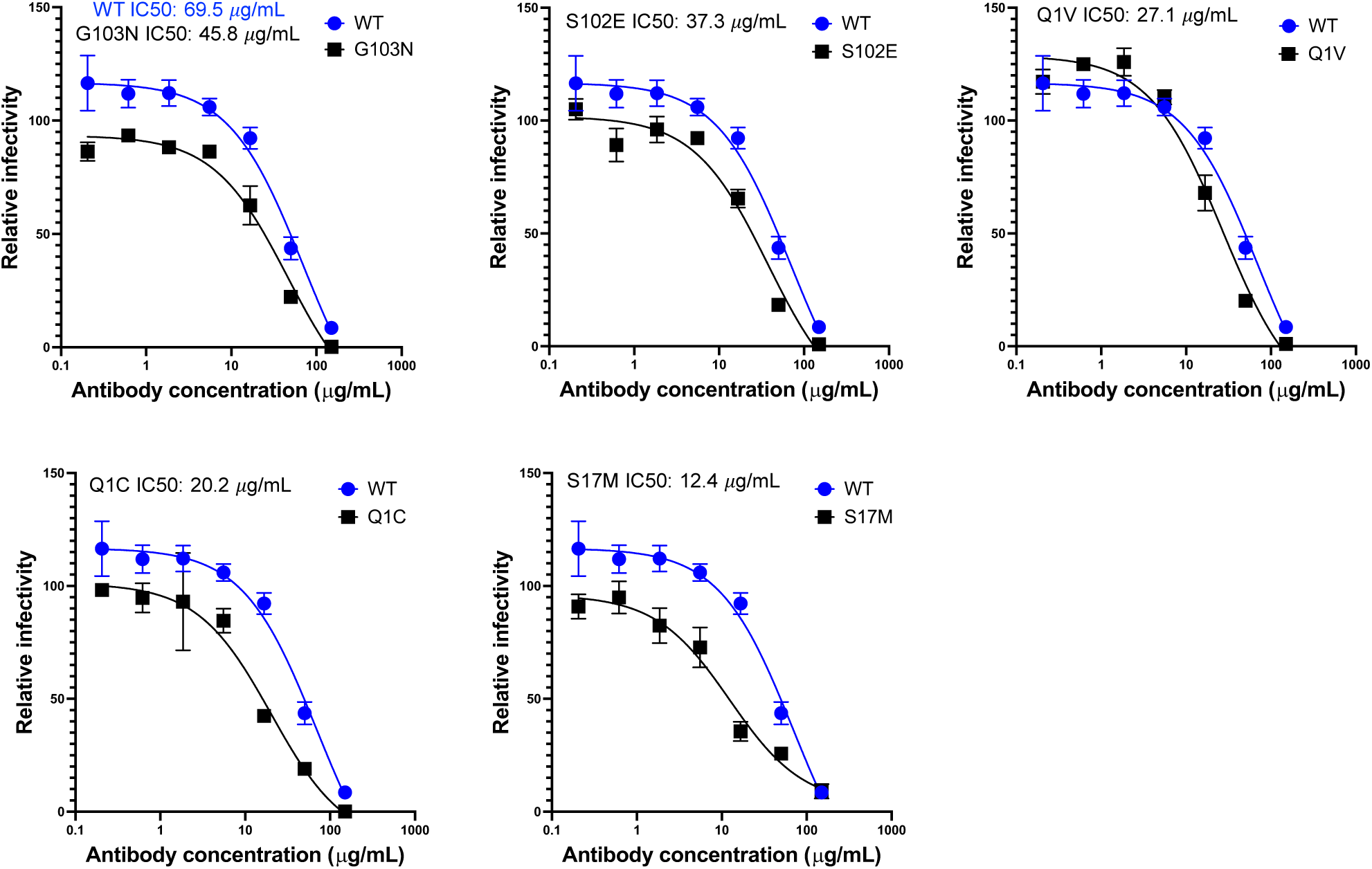
Neutralization results for single mutation variants expressed as IgG1 for validation.

**Figure S4.**
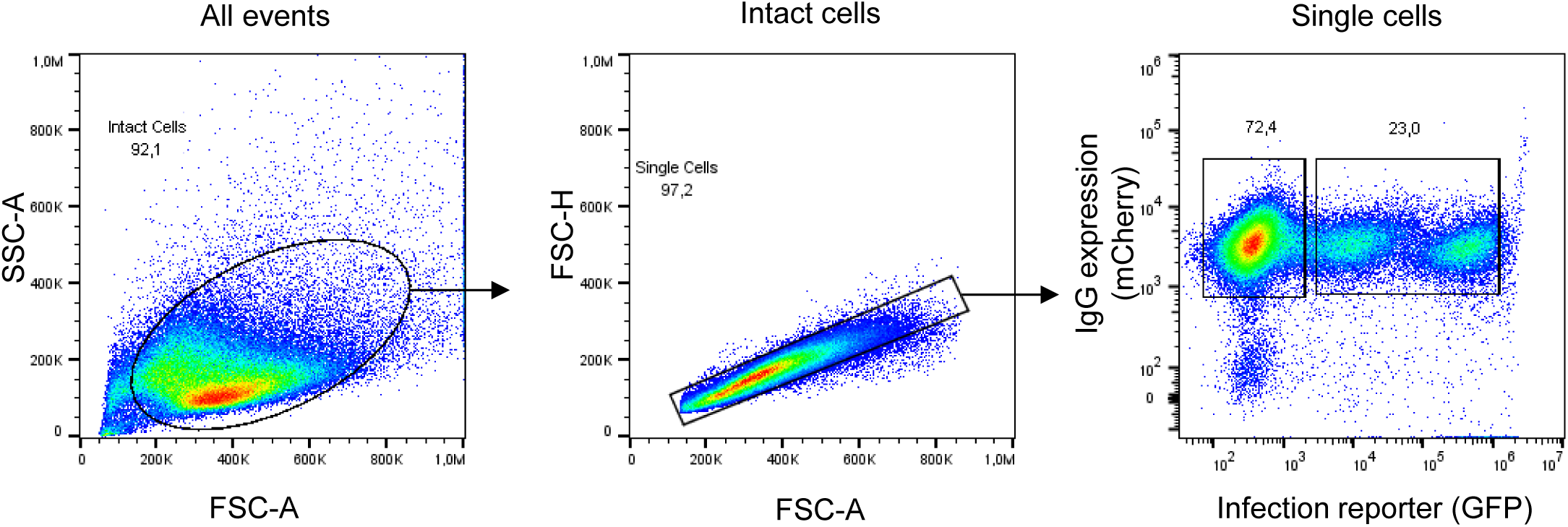
Example flow cytometry gates used in library screening assays.

## Supplementary Tables

**Table S1.** NGS data from oligoclonal library screens. Cell lines expressing a single monoclonal antibody from a panel of antibodies were mixed to generate a synthetic oligoclonal library for screening. Frequency of each clone in a library was calculated from the number of reads for an antibody CDR-H3 in NGS data, divided by the total monoclonal antibody reads in that library.

| A | HIV BG505.W6M.C2<br>in-droplet challenge |  |  |
| --- | --- | --- | --- |
|  | 72A1 | VRC01 | VRC34 |
| IC50 (µg/mL) | >100 | 0.06 | 0.21 |
| Frequency in input library (%) | 90.11 | 3.56 | 6.33 |
| Frequency in mCherry+/GFP-library (%) | 91.56 | 1.49 | 6.94 |
| Frequency in mCherry+/GFP+ library (%) | 99.99 | 0.00419 | 0.00686 |
| Fold-change | 0.916 | 357 | 1,010 |

| B | SARS-CoV-2 D614G<br>in-droplet challenge |  |  |
| --- | --- | --- | --- |
|  | VRC01 | CR3022 | 1-20 |
| IC50 (µg/mL) | >100 | >100 | 0.12 |
| Frequency in input library (%) | 99.01 | 0.399 | 0.589 |
| Frequency in mCherry+/GFP-library (%) | 99.53 | 0.157 | 0.312 |
| Frequency in mCherry+/GFP+ library (%) | 99.64 | 0.298 | 0.0587 |
| Fold-change | 0.998 | 0.529 | 5.31 |

**Table S2.**
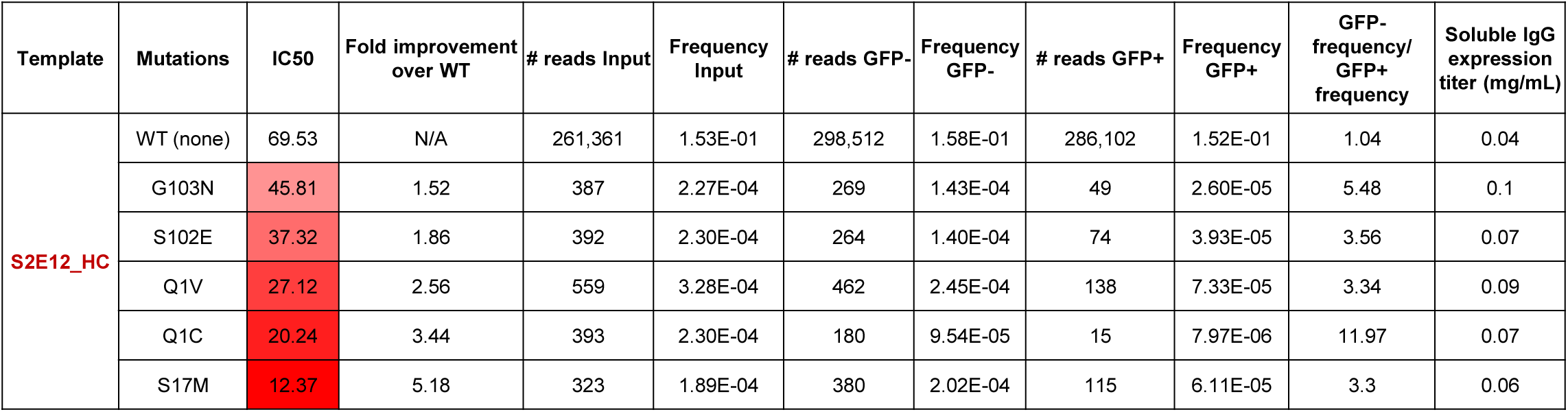
S2E12 clones selected for expression and characterization from heavy chain SSM library screening data.

